# Impacts of Bisphenol A and Ethinyl Estradiol on Male and Female CD-1 Mouse Spleen

**DOI:** 10.1101/109009

**Authors:** Robin B. Gear, Scott M. Belcher

## Abstract

The endocrine disruptor bisphenol A (BPA) and the pharmaceutical 17α-ethinyl estradiol (EE) are synthetic chemicals with estrogen-like activities. Despite ubiquitous human exposure to BPA, and the wide-spread clinical use of EE as oral contraceptive adjuvant, the impact of these estrogenic endocrine disrupting chemicals (EDCs) on the immune system is unclear. Here we report results of *in vivo* dose response studies that analyzed the histology and microstructural changes in the spleen of adult male and female CD-1 mice exposed to 4 to 40,000 μg/kg/day BPA or 0.02 to 2 μg/kg/day EE from conception until 12-14 weeks of age. Results of that analysis indicate that both BPA and EE have dose- and sex-specific impacts on the cellular and microanatomical structures of the spleens that reveal minor alterations in immunomodulatory and hematopoietic functions. These findings support previous studies demonstrating the murine immune system as a sensitive target for estrogens, and that oral exposures to BPA and EE can have estrogen-like immunomodulatory affects in both sexes.

## Introduction

Bisphenol A (BPA) is one of the highest volume chemicals produced worldwide inpart due to its widespread use as a starting material and plasticizer for polymer production ^1^. The global demand for BPA in 2013 was estimated at more than 7 million metric tons, with the demand expected to grow to over 9.6 million metric tons by 2020 ^2^. Biomonitoring studies from the US Centers for Disease Control and Prevention demonstrated that more than 90% of Americans had detectable levels of BPA metabolites in their urine ^3,4^. The majority of human exposure results from ingestion of monomeric BPA contaminants in foods and beverages that results in an estimated mean adult dietary exposure of <0.01-0.40 μg/kg of body weight per day and 0.06 - 1.5 μg/kg/day at the 95^th^ percentile ^1^. Exposure estimates for children range from 0.01 – 13 μg/kg/day ^5^. Because of this wide-spread exposure and the endocrine disruptive actions of BPA, there is much interest in clarifying the possible long-term effects that BPA exposures have on human health and the well-being of animal species.

The disruptive mechanisms of BPA action are related to estrogen signaling, which in some cases can also result in alterations of thyroid hormone and androgen signaling, and abnormal hormonal control of metabolism ^6-8^. Along with concerns related to the effects of estrogenic EDCs like BPA on the reproductive system, there is much scientific and public health interest related to answering the question of whether or not BPA influences the function of the immune system ^9,10^. The idea that environmental estrogens alter immune function is founded upon important differences in the inflammatory responses of females and males. These sexually dimorphic immune responses result from dissimilarities in gonadal steroid hormone levels and the sex-specific immunomodulatory actions of estrogens ^11-13^. It is well established that endogenous estrogens (e.g. 17β-estradiol) can influence all major cell linages of the immune system with resulting consequences on both humoral and cellular immunity ^11,13^. However, the sex-specific immunomodulatory actions of estrogens are complex, as both low and high concentrations of endogenous estrogens have cell-specific and often paradoxical immunomodulatory actions. The complex tissue- and immune-cell specific responses to estrogens are in part, controlled by differential expression of the nuclear hormone estrogen receptors (ER), ERα and ERβ, which regulate cell-specific differences in estrogen-mediated signaling and the resulting differential cellular responses ^12^.

It is also unclear whether, or to what degree, exposures to estrogenic endocrine disrupting compounds can alter or cause harm to immune function. Because of the potential for BPA or EE to disrupt endogenous estrogen signaling via modes of action similar to endogenous estrogens, it is plausible that exposures to each of these environmental estrogens may effect various aspects of the immune response. Numerous *in vivo* and *in vitro* studies have been performed to determine whether BPA, often used experimentally at high concentrations, can impact components of the immune system. For example, effects have been reported on mast cell degranulation, lymphocyte proliferation, antibody responses, modulation of innate immunity, and the function of regulatory T cells ^14–20^. Recently, there has been focused efforts exploring the possible connection between *developmental* exposures to BPA and later inflammatory responses of the lung. In experimental models of influenza infection and asthma, BPA exposures were found to elicit modest effects on innate immunity and very subtle, sex specific effects on airway lymphocytic and lung allergic inflammation ^17,21^.

With the goal of understanding the complex nature of BPA’s effects on a wide range of systems, organs, and end points, we performed studies to determine the physiological impacts of dietary exposures to different doses of BPA, and EE as a control for estrogen-like effects, in CD-1 mice ^22^. During the initial phases of those studies, it was found that both EE and BPA induced uterine pathology that was characterized by markedly increased inflammation and extensive intrauterine macrophage infiltration ^23,24^. The causes of the observed pathogenic increase in estrogen-induced inflammation and the etiology of the resulting uterine pathology remains unresolved. To better understand the nature of the effects of oral exposure to BPA and EE on the observed immunological changes, here we have assessed the histologic and microstructural changes in spleens of male and female CD-1 mice exposed from conception through adulthood to a range of BPA or EE doses ^22^. We focused this study on the spleen, because of its important role in normal reproduction and pathological inflammatory functions in the uterus and ovary. In the uterus and ovary, inflammatory processes are essential for normal estrous (in rodents) and menstrual (in humans) cyclicity, ovulation, implantation, placentation, maternal-fetal tolerance and parturition ^25-28^. The presented results and their analyses add to the available data on the effects of estrogen-like endocrine disrupting pharmaceuticals (EE) and estrogenic environmental contaminants (BPA) to the secondary lymphoid tissue and the hematopoietic compartment of the spleen, and offer foundational insight necessary to establish understanding of how endocrine disruption of the immune system might impact target organ systems.

## Results

The body weights of all animals in the study were monitored from birth until study completion with final body and spleen weights recorded for analysis at time of necropsy (~12 weeks of age). The mean body and spleen weights with standard deviations and sample numbers for each group are presented in Table 1. Two-way ANOVA revealed a significant main effect on body weights of the sex factor *F*(1,104) = 108.9, (p < .0001), and no significant main effect for the BPA exposure factor *F*(5,104) = 1.13, (p = .35). For EE exposure groups, two-way ANOVA also showed a significant main effect on body weights for the sex factor *F*(1,58) = 46.09, (*p* <.0001), with no significant main effect of the exposure factor *F*(3,58) = 2.05 (p = .12).

**Table 1.** Body and Spleen Weights at Necropsy (g)

| | Control | Bisphenol A (ppm) | | | | | 17α-Ethinyl Estradiol (ppm) | | |
| --- | --- | --- | --- | --- | --- | --- | --- | --- | --- |
|  | 0.0 | 0.03 | 0.3 | 3.0 | 30 | 300 | 0.0001 | 0.001 | 0.01 |
| Female Body Weight | 32.61 ± 5.42 (5) | 32.27 ± 4.67 (8) | 32.29 ± 5.77 (9) | 33.85 ± 4.93 (10) | 36.45 ± 3.50 (11) | 34.18 ± 3.67 (13) | 36.55 ± 6.26 (7) | 34.49 ± 3.98 (12) | 31.80 ± 4.54 (6) |
| Male Body Weight | 40.56 ± 6.81 (14) | 44.57 ± 2.66 (10) | 43.23 ± 5.63 (9) | 44.87 ± 3.83 (12) | 42.61 ± 5.32 (7) | 44.34 ± 4.60 (8) | 43.85 ± 3.02 (8) | 44.84 ± 3.93 (6) | 42.30 ± 4.41 (8) |
| Female Spleen Weight | 0.115 ± 0.009 (5) | 0.124 ± 0.014 (8) | 0.128 ± 0.032 (9) | 0.130 ± 0.029 (10) | **0.157 ± 0.049<sup>a</sup> (11)** | 0.133 ± 0.021 (13) | 0.132 ± 0.041 (7) | **0.162 ± 0.035<sup>a</sup> (12)** | 0.150 ± 0.041 (6) |
| Male Spleen Weight | 0.1281 ± 0.0303 (14) | 0.1438 ± 0.0182 (10) | 0.1285 ± 0.0280 (9) | 0.155 ± 0.034 (12) | **0.172 ± 0.046<sup>a</sup> (7)** | 0.137 ± 0.014 (8) | 0.1517 ± 0.023 (8) | 0.142 ± 0.010 (6) | 0.159 ± 0.039 (8) |

Values represent group means ± SD (n); <sup>a</sup> Significantly different from control by Dunnett's multiple comparison test ( $p \leq .05$ )

For spleen weight (Figure 1), two-way ANOVA indicated a significant main effect on spleen weights for both the sex factor *F*(1,102) =4.41, (*p* = .04), and for the BPA exposure factor *F*(5,102) = 5.90, (p < .0001). For EE exposure groups, two-way ANOVA showed no significant main effect on spleen weights of the sex factor *F*(1,58) = 0.83, (*p* = .37), and a significant main effect of the exposure factor *F*(3,58) = 3.49 (p = .02). A one-way analysis of variance indicated significant differences in spleen weights between groups of BPA-exposed males [*F*(5,54) = 2.72, (p = .03)] and females [*F*(5,48) = 3.39, (p = .01)]. Dunnett’s multiple comparison tests indicated that spleen weights for both sexes were significantly different from the mean weight of same sex control spleens in the 30 ppm BPA group, and the mean spleen weight of females in the 0.001 EE group was significantly greater than that of the female control group (Table 1). It is notable that increasing doses of BPA or EE were generally associated with increases in spleen weights with the exception of the highest concentration of BPA (300 ppm), where a significant decrease in spleen weight compared to the 30 ppm exposure group was observed in both females (t(21) = 2.05, p = .03) and males (t(13) = 2.05, p = .04). This finding is suggestive of a complex (i.e. non-monotonic) concentration response relationship between these exposures and spleen weight.

**Figure 1.**
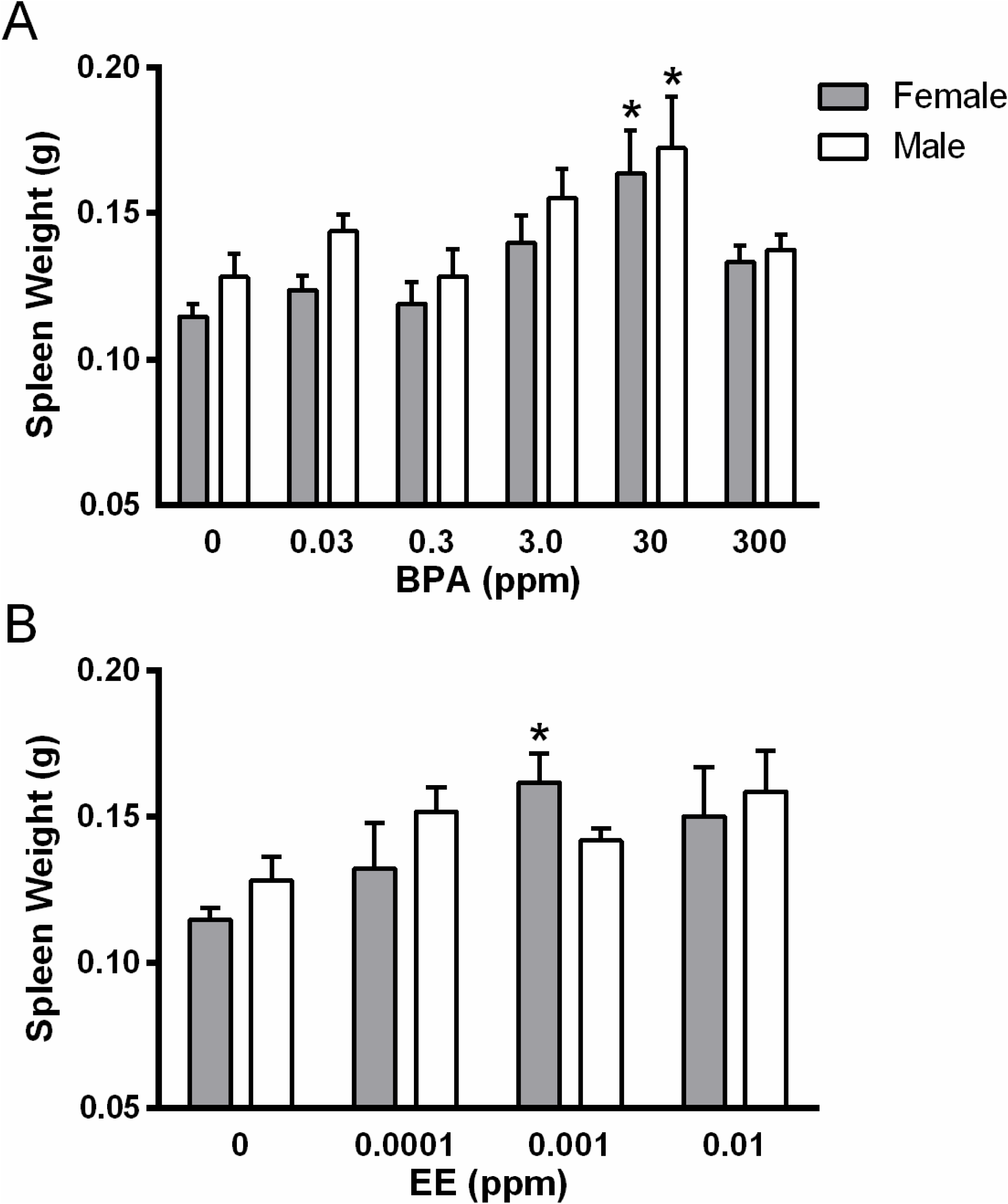
Effects of exposure on spleen weight. Spleen weight of males and females exposed to increasing doses of BPA **(A)** or EE **(B)**. * indicates significantly different from same sex control *p* < .05 by Dunnett’s multiple comparisons test.

## Histopathologic analysis of the Spleen: effects in white pulp

No overtly cytotoxic effects to the spleen lymphoid tissues were appreciated. In general, the nature of the evaluated microscopic alterations were similar for the BPA and EE exposure groups. Compared to unexposed controls, dietary exposure to BPA or EE generally resulted in minimal decreases in the size, numbers, and cellularity of the periarteriolar lymphoid sheaths (PALS) with decreased numbers of primary lymphoid follicles (Figures 2 and 3). With the exception of the male 3 ppm exposure group, there was no significant effects on germinal centers (secondary lymphoid follicles), and modest effects on the B-cell rich marginal zones (Table 2).

**Figure 2.**
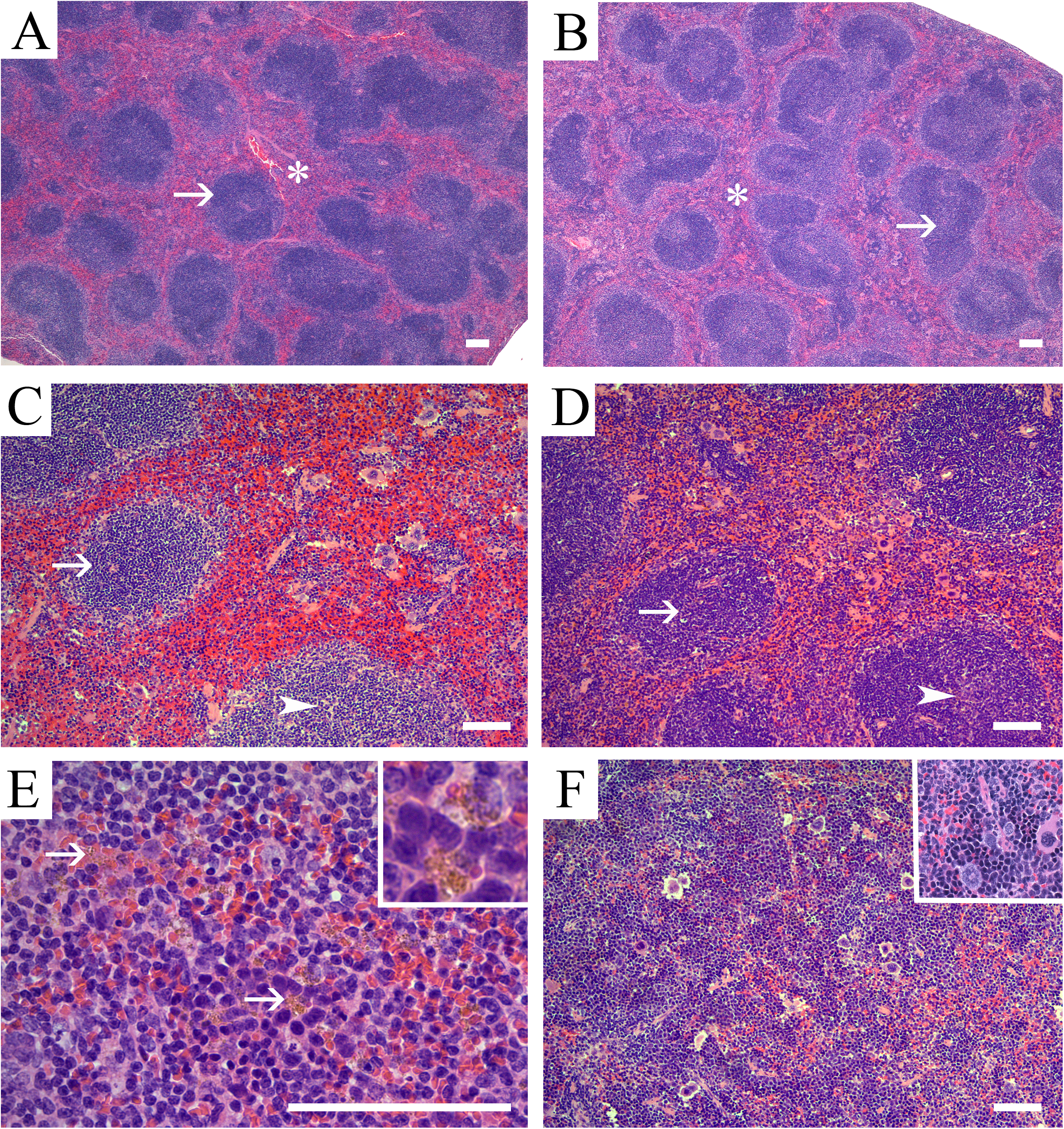
*Representative photomicrographs of the splenic microanatomical structures in Control and* 17α-ethinyl estradiol *exposed adult male and female CD-1 mice.* Shown are representative H&E stained sections of splenic tissue from **(A)** control male and **(B)** control female mice illustrating the normal micro-architecture of the white pulp (arrow) and red pulp (asterisk). Note that control tissue displays abundant white pulp components. **(C)** Representative images of males exposed to dietary 17α-ethinyl estradiol (0.01 ppm EE). Decreased periarteriolar lymphoid sheath (PALS) numbers, size and cellularity (arrow), together with paucicellular primary lymphoid follicles (arrowhead) are evident. **(D)** Shown is an image for a representative female from the 0.01 ppm EE exposure group showing decreased PALS numbers and cellularity (arrow) with discrete primary lymphoid follicles and mildly decreased numbers of follicular lymphocytes (arrowhead). **(E)** Representative image of an EE-exposed male from the 0.01 ppm exposure group which demonstrates increased amounts of golden-yellow pigment (hemosiderin) in the red pulp that was often contained within macrophages (arrows and inset) **(F)** Representative image from a 0.0001 ppm EE exposed female demonstrating the observed phenotype of diffuse expansion of the red pulp by extramedullary hematopoiesis (EMH). The EMH was characterized by the presence of myeloid, megakaryocytic precursors, and a morphological predominance of erythroid precursors (inset). Scale bars indicate 100 μm.

**Figure 3.**
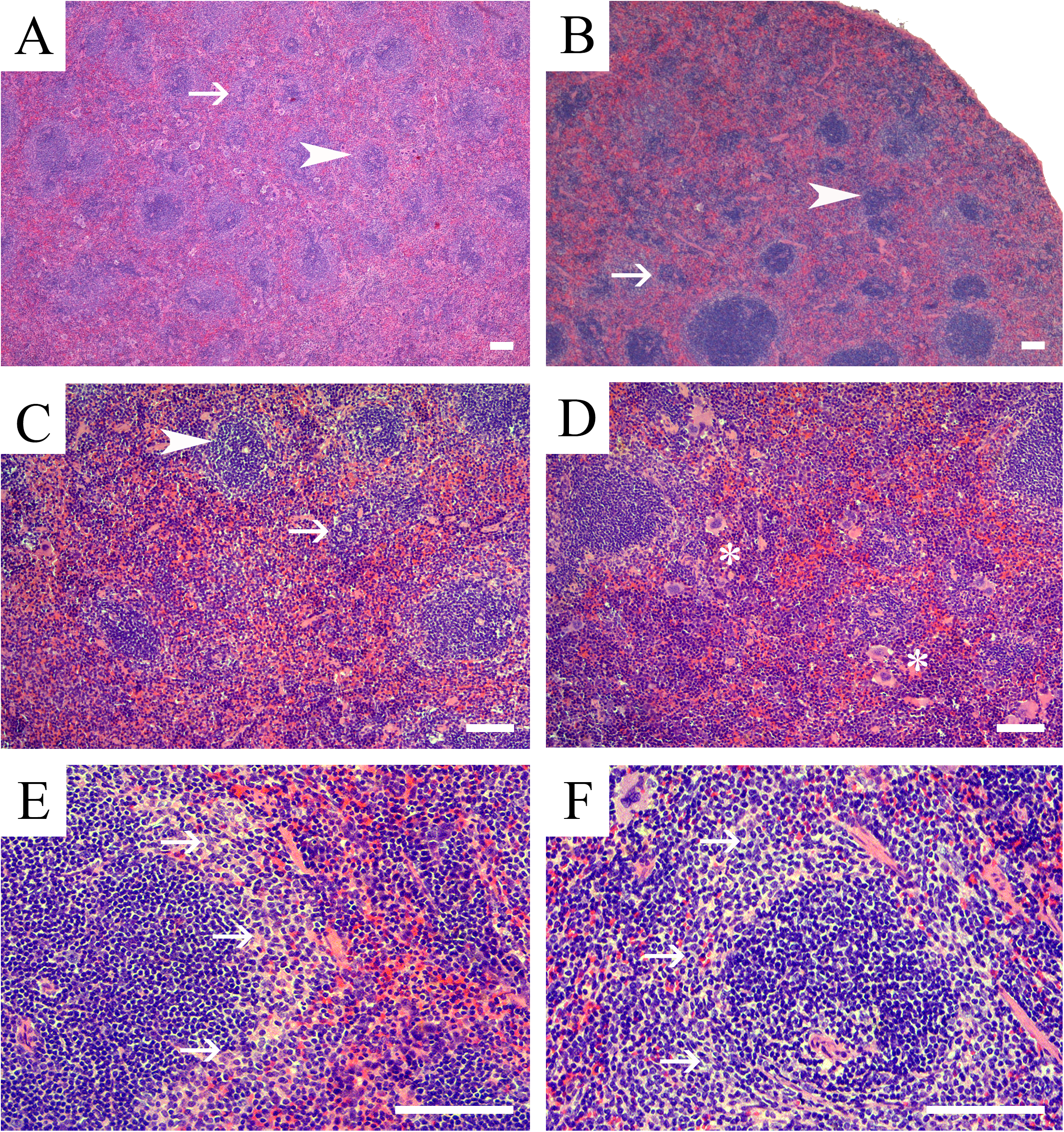
*Representative photomicrographs of splenic microanatomical structures in EE and BPA-exposed adult male and female CD-1 mice*. Shown are representative H&E stained sections of splenic tissue from **(A)** a male mouse exposed to BPA (0.03 ppm), and **(B)** a female mouse exposed to 3 ppm BPA. These are illustrative of the representative phenotypes in males and females that was characterized by decreased PALS (arrowhead) numbers, size, and lymphocyte cellularity (arrow). These effects were typically more pronounced and diffuse in male versus females. **(C)** High magnification image of a male spleen representative of the 0.03 ppm BPA exposure group and demonstrating decreased PALS (arrowhead) size, numbers and lymphocyte cellularity (arrow), along with decreased primary lymphoid follicles cellularity (arrow). **(D)** High magnification image of a representative female spleen from the 3 ppm BPA exposure group characterized by decreased PALS numbers and cellularity with decreased lymphoid follicle cellularity. In addition to the white pulp changes, BPA-exposed females also had increased hematopoietic precursor populations (EMH) within the red pulp, which were predominantly of erythroid lineage (asterisks). Shown is a representative image of a section from a 0.0001 ppm EE-exposed male **(E)** and female from the 0.03 BPA group **(F)** showing prominence of marginal zones, a phenotype that was observed in primarily males from EE exposure groups and female from both BPA- and EE-groups. Scale bars indicate 100 μm.

**Table 2.** Histopathology of Spleen White Pulp

|  |  |  | PALS |  |  |  |  |  | Marginal zone |  | Follicles |  |  |  |  |  |
| --- | --- | --- | --- | --- | --- | --- | --- | --- | --- | --- | --- | --- | --- | --- | --- | --- |
|  |  |  | size |  | numbers |  | lymphocytes |  | size |  | numbers |  | lymphocytes |  | germinal centers |  |
| *BPA (ppm)* | Sex | n= | AFF | UN | AFF | UN | AFF | UN | AFF | UN | AFF | UN | AFF | UN | AFF | UN |
| 0 | F | 5 | 0 | 5 | 0 | 5 | 0 | 5 | 0 | 5 | 0 | 5 | 0 | 5 | 0 | 5 |
| 0.03 | F | 8 | 4 | 4 | 5 | 3 | 5 | 3 | 2 | 6 | 0 | 5 | 6 | 2 | 0 | 8 |
| 0.3 | F | 9 | 0 | 9 | 3 | 6 | 4 | 5 | 4 | 5 | 4 | 5 | 5 | 4 | 1 | 8 |
| 3 | F | 10 | 0 | 5 | 6 | 4 | 6 | 4 | 2 | 8 | 3 | 7 | 2 | 8 | 1 | 9 |
| 30 | F | 11 | 5 | 6 | 5 | 6 | 6 | 5 | 5 | 6 | 4 | 7 | 5 | 6 | 1 | 10 |
| 300 | F | 13 | 6 | 7 | 5 | 8 | 7 | 6 | 5 | 8 | 5 | 8 | 6 | 7 | 0 | 13 |
|  |  |  | size |  | numbers |  | lymphocytes |  | size |  | numbers |  | lymphocytes |  | germinal centers |  |
| *EE (ppm)* | Sex | n= | AFF | UN | AFF | UN | AFF | UN | AFF | UN | AFF | UN | AFF | UN | AFF | UN |
| 0.0001 | F | 7 | 2 | 5 | 4 | 3 | 4 | 3 | 3 | 4 | 3 | 4 | 2 | 5 | 0 | 7 |
| 0.001 | F | 12 | 5 | 7 | 9 | 3 | 8 | 4 | 4 | 8 | 2 | 10 | 4 | 8 | 0 | 12 |
| 0.01 | F | 6 | 5 | 4 | 3 | 3 | 5 | 1 | 2 | 4 | 1 | 5 | 0 | 6 | 0 | 6 |
|  |  |  | size |  | numbers |  | lymphocytes |  | size |  | numbers |  | lymphocytes |  | germinal centers |  |
| *BPA (ppm)* | Sex | n= | AFF | UN | AFF | UN | AFF | UN | AFF | UN | AFF | UN | AFF | UN | AFF | UN |
| 0 | M | 14 | 0 | 14 | 0 | 14 | 0 | 14 | 0 | 14 | 0 | 14 | 0 | 14 | 0 | 14 |
| 0.03 | M | 7 | 0 | 7 | 5 | 5 | 4 | 6 | 0 | 7 | 3 | 7 | 3 | 7 | 0 | 10 |
| 0.3 | M | 9 | 6 | 3 | 6 | 3 | 5 | 4 | 0 | 8 | 3 | 6 | 4 | 5 | 1 | 8 |
| 3 | M | 12 | 4 | 8 | 3 | 9 | 4 | 8 | 6 | 6 | 6 | 6 | 4 | 8 | 0 | 12 |
| 30 | M | 7 | 3 | 4 | 2 | 5 | 5 | 2 | 2 | 5 | 4 | 3 | 2 | 5 | 3 | 4 |
| 300 | M | 8 | 2 | 6 | 2 | 6 | 5 | 3 | 0 | 8 | 1 | 7 | 1 | 7 | 0 | 8 |
|  |  |  | size |  | numbers |  | lymphocytes |  | size |  | numbers |  | lymphocytes |  | germinal centers |  |
| *EE (ppm)* | Sex | n= | AFF | UN | AFF | UN | AFF | UN | AFF | UN | AFF | UN | AFF | UN | AFF | UN |
| 0.0001 | M | 8 | 3 | 5 | 6 | 2 | 5 | 3 | 2 | 6 | 4 | 4 | 4 | 4 | 2 | 6 |
| 0.001 | M | 6 | 3 | 3 | 1 | 5 | 3 | 3 | 2 | 4 | 2 | 4 | 2 | 4 | 1 | 5 |
| 0.01 | M | 8 | 3 | 5 | 2 | 6 | 3 | 5 | 2 | 6 | 3 | 5 | 2 | 6 | 1 | 7 |

Values are number of samples affected or unaffected. AFF: indicates affected; UN: indicates unaffected. Significant differences compared to same sex control are indicated in bold with gray shading.

### Effects of exposures in white pulp of females

The observed effects resulting from BPA or EE exposures on the white pulp of females included changes in PALS number and size (Figure 2B; 2D; 3B). The histopathologic phenotypes observed in the T-cell rich PALS region of affected females were predominantly minimal to mild decreases in size, number, and cellularity of the PALS, with minimal to mild increases in apoptotic lymphocytes. With the exception of changes in PALS size at the lowest EE dose (χ^2^ (1, *n* = 12) = 1.7, *p* = .10), significant effects of EE exposure on incidence of changes in PALS number, size and cellularity changes were observed for each endpoint in females (Table 2). PALS size in the female 0.3 BPA group was unchanged compared to control, and while decreases in PALS numbers were evident, the incidence was also not significantly different from control (χ^2^ (1, *n* = 14) = 2.1, *p* = .07), although a significant decrease (χ^2^ (1, *n* = 14) = 3.1, *p* = .04) in cellularity was observed in that exposure group.

Effects on lymphocyte numbers also varied depending on exposure group with a predominance of decreased lymphocyte numbers resulting from BPA or EE exposure. Changes in the size of the marginal zone were also noted for females from each exposure group, with the highest percentage (46%) of affected individuals observed in the 30 ppm BPA group ((χ^2^ (1, *n* = 16) = 3.3, *p* = .03); Table 2). In each of the EE exposure groups, females presented with mild increases in their marginal zone size (Figure 3F), the incidence of these changes were not significant. A general decrease in the number of follicles was also noted, with a significant decrease in lymphocyte numbers present in the follicles observed in each BPA exposure group except the 3 BPA group (χ^2^ (1, *n* = 15) = 1.15, *p* = .14). No significant effects on follicles was observed in any EE exposure group and the numbers of germinal centers of BPA or EE exposed females were not significantly altered (Table 2).

### Effects of exposures in white pulp of males

In males both EE (Figure 2C) and BPA (Figures 3A; 3C) exposures resulted in significant changes in severity of altered PALS numbers (*H* = 13.9, *p* = .02), PALS size (*H* = 11.8, *p* = .03), and the numbers of follicles (*H* = 11.7, *p* = .03) and lymphocytes (*H* = 12.8, *p* = .02). These effects were predominantly minimal to mild decreases in PALS size and number, with changes of moderate severity grade observed at the lowest dose of BPA for effects on PALS size, number, and cellularity. There was no change in PALS size observed in the lowest BPA dose group and PALS numbers at 0.001 EE (χ^2^ (1, *n* = 20) = 2.5, *p* = .06) was not significantly different from control. Significant effects on PALS size, numbers and lymphocyte numbers were detected in all other exposure groups (Table 2). Significant increases in size of the marginal zones were noted in all EE exposure groups (Figure 3E) and in the 3 ppm (χ^2^ (1, *n* = 22) = 9.1, *p* = .001) and 30 ppm (χ^2^ (1, *n* = 21) = 4.4, *p* = .02) BPA groups. Significant increases in the numbers of follicles were observed for all treatment groups except for the 300 ppm BPA group ((χ^2^ (1, *n* = 22) = 1.83, *p* = .09), Table 2). The numbers of lymphocytes localized to follicles were significantly decreased for all treatment groups except for the 300 ppm BPA group (χ^2^ (1, *n* = 22) = 1.83, *p* = .09). Germinal center numbers were more frequently altered in males, where significant increases were observed in the 0.001 EE (χ^2^ (1, *n* = 22) = 3.9, *p* = .03) and 30 BPA (χ^2^ (1, *n* = 21) = 7.0, *p* = .004) exposure groups (Table 2).

## Histopathologic analysis of the Spleen: effects in the red pulp

Overall, the general effects of treatment on the red pulp were similar in males and females (Table 3). Observed effects consisting primarily of minimal to mild (BPA) and minimal to moderate (EE) increases in cellularity resulting from increased hemosiderin pigment (Figure 2E) and extramedullary hematopoesis (EMH; Figure 2F). A higher percentage of females across all diets had increased EMH when compared to exposed males with the exception of the highest dose groups. Observed increases in EMH were characterized by a predominance of erythroid precursors (monolineage) in all affected animals (Table 3; Figure 2F). The second most commonly increased population was the megakaryocytic precursor lineage. The severity grade for increased hemosiderin pigment in BPA-exposed females (*H* = 11.6, *p* = .04) and males (*H* = 11.3, *p* < .05) was mostly minimal, with mild increases observed in the highest exposure groups. In contrast, the effect in EE-exposed females (*H* = 12.72, *p* = .005) was mainly minimal with moderate increases in hemosiderin pigment observed only at the lowest EE dose. Increases in hemosiderin pigment in EE-exposed males was observed at the two highest doses and ranged from minimal to mild in severity (*H* = 13.13, *p* = .004).

**Table 3.** Histopathology of Spleen Red Pulp, Tingible-Body Macrophages and Hemosiderin

|  |  |  | Red Pulp |  |  |  | Hemosiderin |  | Tingible-Body Macrophages |  |  |  |  |  |
| --- | --- | --- | --- | --- | --- | --- | --- | --- | --- | --- | --- | --- | --- | --- |
|  |  |  | size |  | hematopoietic cells |  | pigment |  | germinal centers |  | PALS |  | red pulp |  |
| *BPA (ppm)* | Sex | n= | AFF | UN | AFF | UN | AFF | UN | AFF | UN | AFF | UN | AFF | UN |
| 0 | F | 5 | 0 | 5 | 0 | 5 | 0 | 5 | 0 | 5 | 0 | 5 | 0 | 5 |
| 0.03 | F | 8 | **6** | **2** | **6** | **2** | 1 | 7 | 0 | 8 | 3 | 5 | 2 | 6 |
| 0.3 | F | 9 | 4 | 5 | 4 | 5 | **6** | **3** | 0 | 9 | 2 | 7 | 0 | 9 |
| 3 | F | 10 | 4 | 6 | 7 | 3 | **5** | **5** | 0 | 10 | 1 | 9 | 1 | 9 |
| 30 | F | 11 | **6** | **5** | **9** | **2** | 4 | 7 | 0 | 11 | **5** | **6** | 2 | 10 |
| 300 | F | 13 | 5 | 8 | **6** | **7** | **9** | **4** | 0 | 13 | 2 | 11 | 0 | 13 |
|  |  |  | size |  | hematopoietic cells |  | pigment |  | germinal centers |  | PALS |  | red pulp |  |
| *EE (ppm)* | Sex | n= | AFF | UN | AFF | 0 | AFF | 0 | AFF | 0 | AFF | 0 | AFF | 0 |
| 0.0001 | F | 7 | 5 | 2 | **7** | **0** | **7** | **0** | 0 | 7 | 1 | 6 | 0 | 7 |
| 0.001 | F | 12 | 7 | 5 | **12** | **0** | 5 | 8 | 0 | 12 | 0 | 12 | 0 | 12 |
| 0.01 | F | 6 | **4** | **2** | **4** | **2** | **5** | **1** | 0 | 6 | 1 | 5 | 0 | 6 |
|  |  |  | size |  | hematopoietic cells |  | pigment |  | germinal centers |  | PALS |  | red pulp |  |
| *BPA (ppm)* | Sex | n= | AFF | UN | AFF | UN | AFF | UN | AFF | UN | AFF | UN | AFF | UN |
| 0 | M | 14 | 0 | 14 | 0 | 14 | 0/0 | 14 | 0 | 14 | 0 | 14 | 0 | 14 |
| 0.03 | M | 7 | **2** | **8** | **2** | **8** | **2** | **8** | 0 | 10 | 0 | 10 | **2** | **8** |
| 0.3 | M | 9 | **6** | **3** | **3** | **6** | 1 | 8 | 0 | 9 | 2 | 7 | 0 | 9 |
| 3 | M | 12 | **7** | **5** | **6** | **6** | 2 | 10 | 0 | 12 | 1 | 11 | 0 | 12 |
| 30 | M | 7 | **5** | **2** | **3** | **4** | **3** | **4** | 1 | 6 | 1 | 6 | 0 | 7 |
| 300 | M | 8 | 0 | 8 | **6** | **2** | **4** | **4** | 0 | 8 | 0 | 8 | 0 | 8 |
|  |  |  | size |  | hematopoietic cells |  | pigment |  | germinal centers |  | PALS |  | red pulp |  |
| *EE (ppm)* | Sex | n= | AFF | UN | AFF | 0 | AFF | 0 | AFF | 0 | AFF | 0 | AFF | 0 |
| 0.0001 | M | 8 | **2** | **6** | 3 | 5 | 0 | 8 | 0 | 8 | **2** | **6** | 0 | 8 |
| 0.001 | M | 6 | **3** | **3** | 4 | 2 | 3 | 3 | 0 | 6 | 0 | 6 | 0 | 6 |
| 0.01 | M | 8 | **2** | **6** | 7 | 1 | 4 | 4 | 0 | 8 | 1 | 7 | 0 | 8 |

Values are number of samples affected or unaffected. AFF: indicates affected; UN: indicates unaffected.
Significant differences compared to same sex control are indicated in bold with gray shading.

## Discussion

Here we have reported results from *in vivo* dose response studies that analyzed the histological and microstructural changes in the spleen of male and female CD-1 mice exposed to BPA or EE from conception until 12-14 weeks of age ^22^. Based on our previous analysis that found BPA and EE increased uterine macrophage numbers and inflammation ^23,24^, the aim of this study was to investigate further the estrogen-like inflammatory responses induced by oral exposure to these estrogenic chemicals. The findings and interpretations of this analysis confirmed that both BPA and EE have dose and sex specific effects in the spleen consistent with alterations in immunomodulatory and hematopoietic functions. These effects were modest with minimal to moderate changes observed in the different microanatomical structures of spleens of CD-1 mice as a result of oral exposures to BPA from 4 μg/kg/day to 40 mg/kg/day, or to 0.02 to 2 μg/kg/day of the orally bioavaliable estrogenic drug EE. Those findings support previous studies that have defined the murine immune system as a sensitive target for the endogenous estrogen hormone 17β-estradiol ^29^. Further, the detection of these influences in the spleen also supports our previous findings that demonstrated BPA and EE each have immunomodulatory impacts on both sexes of the CD-1 mouse ^23,24^.

The foundations of the presented studies were based on our previous observations that the estrogenic actions of BPA and EE could induce pathologic fibrosis and inflammatory pathology of the uterus of exposed females ^23,24^. Other previous studies have investigated the potential for the endocrine disruptor BPA to disrupt immune function by focusing on determining whether BPA exposure altered various physiologic properties of isolated immune cells, by establishing an association between exposure and inflammatory or autoimmune disease, or whether exposure to BPA modified immune or inflammatory responses in specific animal models ^9,10,30^.

Because the impacts of estrogens, and by extension estrogenic endocrine disruptors, are influenced by a variety of factors, the studies presented here took a general approach by assessing whether BPA had observable effects on the spleen. That approach allows for a detailed study-wide approach geared toward developing integrated understanding of the organ-specific immunomodulatry effects of BPA, which can vary depending on animal model, the level and timing of exposures, specific disease model employed, as well as the specifics of disease etiology and progression in different target tissues ^13,30,31^.

The spleen, as a specialized lymphoid organ, is located within the circulatory system and is positioned to monitor and respond to hormonal and environmental factors including infections. Because of its critical role in the inflammatory response, the spleen is also expected to be a sensitive sentinel organ for detecting immunomodulatory influences of environmental factors such as chemical pollutants or pharmaceuticals that impact normal hormone-regulation of immune responsiveness. As a primary indicator of immunotoxicty or altered immunomodulatory actions of the estrogenic compounds BPA and EE in the spleen, we chose to employ the enhanced histopathology (EH) approach as described and adopted in the best practice guidelines for pathological examination of the immune system as developed by the Society of Toxicologic Pathology Immunotoxicology Workforce ^32,33^. The EH approach is especially useful for identifying effects on the immune system and at the same time aids in the identification of the putative target cell population affected, this approach however does not elucidate functional endpoints or define harmful effects ^34^. As a secondary check of the EH findings, and as an aid in developing dose and region-specific interpretations of the EH data, we used an “affected method” as a quantitative incidence analysis to augment the EH approach ^35,36^. Because the nominal nature of the resulting data this approach generally lacks comparative power (sensitivity) for statistically detecting modest differences between control and exposed groups ^35^. The limitations in sensitivity were most pronounced in females due to the smaller number of available controls samples, while there was sufficient power to detect alterations, the lack of detecting significant effects in some groups should be interpreted cautiously especially as it relates to identification of nonmontonic dose responses. While there are limitations, the ability to quantitatively demonstrate deviations from the normal control range of phenotypes was considered to add valuable insight and confidence for interpretation of the dose specific impacts of each compound.

The choice of an appropriately sensitive animal model and strain has proven a critical factor for generating informative data when assessing the immunomodulatory actions of endocrine disruptors in a specific organ or cell type ^23,24,31^. Our Initial evidence pointed toward BPA exposures resulting in tissue-specific estrogen-like increases in inflammation in both C57BL/6J and CD-1 strains of mice ^23,24^. Results of those studies found that C57BL/6J mice were 100 to 1000-fold more sensitive to some estrogen-like actions of BPA and EE then the CD-1 mouse.

However fecundity in the C57BL/6J mouse was decreased by exposure to both BPA and EE across the dose range of interest, which limited the utility of this strain for these dose response analysis ^24^. Whereas the C57BL/6J may serve as a model for detecting disruptive effects in sensitive sub-populations, outbred male and female CD-1 mice are also sensitive to BPA exposure during development and unlike the C57BL/6J strain, do not have any particular inflammatory phenotype propensity. Therefore the CD-1 strain was considered an appropriate model for assessing whether these estrogen endocrine disruptors had detectable immunomodulatory effects on the spleen. The sensitivity of the CD-1 mice to oral estrogenic compounds at the analyzed endpoints was demonstrated here by including the concurrent analysis of the effects of EE which served as a control for detection of estrogen-like effects. The differences in immune-sensitivity to the effects of BPA between the C57BL/6 and CD-1 strains suggest there may exist important gene by environmental differences that influence susceptibility to the immunomodulatory impacts of environmental estrogens. If human sensitivity to estrogenic EDCs are similarly variable, which is considered likely, the findings of effects on some endpoints at the lowest dose analyzed here (4 μg/kg/day) suggest that sensitive individuals may be impacted by exposures to BPA that are currently being experienced in most human populations^1,9^. Additional experimental studies employing genetic models such as diversity outcross or collaborative cross mouse models, will be required to assess the variability of sensitivity to environmental estrogens and define factors that influence these differential impacts on the immune system ^37-39^.

Along with considerations related to individual and population sensitivity, the known sexually-dimorphic nature of immune responses also dictated that immunotoxicity screening studies are able to differentiate sex-specific immunomodulatory effects of endocrine disruptors. As a result, all endpoints analyzed here were considered separately for males and females based on known differential responses and sensitivity. Impacts of exposure were found in both males and females, with sex specific effects of BPA and EE exposure observed in both the PALS and follicles of the white pulp of males and females. The white pulp of the spleen is composed from the small branches of the splenic arteries which are sheathed with lymphoid tissues and organized into the T-cell rich PALS regions, and B-cell rich follicle regions. Effects were generally consistent with the known anti-inflammatory or immunosuppressive influences of exogenous estrogenic compounds at pharmacological concentrations and findings from numerous studies that found minor effects of BPA on both T-and B-cells ^9,11,40,41^.

The red pulp compartment constitutes the venous system of the spleen that is composed from a subset of the smallest branches of the splenic arteries that extend across the B-cell rich marginal zone of the white pulp. Functionally, the red pulp is where senescent or damaged erythrocytes, cellular debris, and infectious microorganisms are remove from circulation and where iron is recycled ^42^. The subcapsular red pulp also acts as an important reservoir for monocytes that can be rapidly mobilization to populate inflammatory sites such as the uterus ^43^. Exposure to either BPA or EE resulted in alterations of the red pulp that consisted chiefly of increased cellularity due to increased EMH that was associated with increased hemosiderin pigment. In mice the spleen is an active hematopoietic organ throughout life, and observed changes in EMH were remarkable only when they exceeded the range for normal in the control spleens. The highest severity grade for these changes was observed in the EE-exposed animals with increased EMH characterized by a predominance of erythroid precursors (monolineage), with the second most commonly increased population being the megakaryocytic precursor lineage. This finding was similar to what was previously described for estrogen-induced myelotoxicity in canines ^44^ and experimentally induced erythropoiesis in mice resulting from estradiol benzoate induced suppression of bone marrow hematopoiesis ^45^. In teleost fish exposure to EE or the estrogenic EDC nonylphenol has been found to cause anemia, leukopenia, and a resulting compensatory multisystemic EMH ^44-46^. Together those finding suggest that the splenic EMH observed here in response to EE and BPA may also be a compensatory response to estrogen-like suppression of bone marrow hematopoiesis. Additionally, E2 exposures that suppress hematopoiesis also result in decreases in peripheral erythrocytes and hemoglobin concentrations that is in part due to increased lysis of erythrocytes ^45,47^. Hemosiderin is a breakdown product of hemoglobin metabolism and recycling (i.e. iron cycle), and its accumulation in the spleen reflects increased lysis of erythrocytes ^42,45,47^. It is possible that the observed increases in hemosiderin found in response to increasing doses of BPA and EE also reflects increased iron cycling due to erythrocyte lysis which may further necessitate increased compensatory EMH. These observations suggest that BPA and EE may have complex effects on hematopoiesis, and erythrocyte turnover or the process of iron-recycling. Additional experimental clarification will be needed to delineate the potential role of these mechanisms in mediating the phenotypic alterations of the spleen resulting from BPA and EE exposure.

Sex-specific differences were evident for a subset of the splenic microstructural endpoints for both BPA and EE. Those effects were also generally consistent with the known sexual dimorphism and differential estrogen-sensitivity of the mammalian immune system’s structure and function ^29,48^. For a given dose of BPA or EE, a higher proportion of treated females displayed microstructural changes in the splenic white pulp compartment with dose-dependent increases in effects notable for most endpoints. With the exception of the highest doses of BPA or EE, in red pulp greater percentages of females also presented with increased EMH compared to males from the same exposure groups. While statistical analysis of ordinal data limits sensitivity of detecting significant dose response relationships across sexes for all endpoints, some provisional inference is possible. In males and females, the effects of EE generally increased with dose reflecting the anticipated increase in estrogen-like activity in each sex. It is notable that the highest frequency of increased EMH in females was at the lowest dose with effects decreasing with increasing dose, whereas a dose-dependent increase in EMH was observed in males exposed to EE. By contrast, for endpoints affected by BPA in white pulp of males, significant effects were often observed in the lowest BPA dose groups, with notable decreases in effects observed with increasing dose. The effects of BPA in the white pulp compartment of females were somewhat more complex. The frequency of observed effects on PALS size was about 50% at the lowest and two highest doses of BPA, with a remarkable lack of effect seen in the middle dose range.

While somewhat complicated by a lack of sensitivity, all of the sex- and concentration-specific effects observed are consistent with phenotypes resulting from both EE and BPA having agonist or partial agonist estrogen-like actions with resulting phenotypes dependent upon the relatively high levels of endogenous estrogens in females and the lower levels of circulating estrogen typical of males. Previous studies have also demonstrated that rapid nongenomic effects of BPA are independent of the nuclear receptor activity of the ERs at very low doses of BPA and are mechanistically responsible for a variety of nonmonotonic dose responses to endogenous estrogens and EDCs ^6, 49-52^. Thus it is possible that the differential cell and sex specific dose response relationships noted involve the integrated actions of rapid signaling at low exposures and increasing nuclear receptor actions at higher exposures of BPA.

Understanding the varied and often paradoxical interactions between endogenous sex hormones, EDCs and the immune system remains challenging. As a result of generally higher levels of circulating and local estrogens, the immune response of females to foreign and/or self-antigens generally results in heightened inflammatory responses with beneficial increases in resistance to infection compared to males. However, this enhancement of immune responsiveness is also responsible for an increased prevalence of harmful autoimmune pathology in females ^11-13^. Paradoxically, it is also well established that low plasma levels of E2, for example during estrus in female rodents or in postmenopausal women, can induce pro-inflammatory phenotypes characterized by Th1-skewing and expansion of IL-1, TNF-α, and IL-6 secreting T-cells in bone marrow ^53^. Whereas periods of high physiological levels of E2, such as proestrus in female rodents, during gestation, or the periovulatory stage of the menstrual cycle in humans, are characterized by an anti-inflammatory phenotype ^11-13^. Similar anti-inflammatory effects are also observed in response to pharmacological concentrations of estrogens in experimental animal models and in humans. A predictive mechanistic understanding the immunomodulatory impacts of exogenous estrogenic compounds, whether induced by pharmaceutical endocrine disruptors such as EE, or from exposures to environmental estrogenic compounds (e.g. dietary phytoestrogens or xenoestrogen contaminants such as BPA) is especially challenging. Along with the fundamental complexity of the inflammatory response to differing concentrations of these estrogen-like compounds being layered upon fluctuating levels of circulating and local hormone levels, predicting the effects of exogenous estrogens is complicated further because each compound has unique, and most often poorly understood impacts on endogenous estrogen responses due to their varying ability to act as estrogen mimics, partial agonists, and/or antagonists and their activity at different receptor systems. The findings of this study support and extend previous findings in experimental animals, and an increasing body of humans studies that suggest links between BPA exposures and varied impacts on immune responsiveness, autoimmunity, and allergic diseases ^14-20^, ^54-58^.

## Methods

### Animal husbandry and exposures

All animal procedures were performed in accordance with protocols approved by the University of Cincinnati Institutional Animal Care and Use Committee and followed the recommendations of the Panel on Euthanasia of the American Veterinary Medical Association. All animal procedures including study specific details of husbandry, breeding and exposures followed the recommendations for best practices for the study of BPA’s endocrine disruptor activity in rodents ^55^. The experimental design, approaches and other experimental details of the study were presented in detail elsewhere ^22^. Animals were maintained on a 14 hr light, 10 hr dark cycle with diets and drinking water provided *ad libitum*. Study animals were housed in single-use polyethylene cages that are certified BPA-free (Innovive, San Diego, CA) with Sani-chip bedding (Irradiated Aspen Sani-chip; P.J. Murphy Forest Products Corp. Montville, NJ) to avoid mycoestrogen-contaminated corncob bedding ^55,56^. Sterile drinking water with oxidizable organic chemicals reduced to <1% of source (<5 ppb) was generated by a dedicated water purification system (Millipore Rios 16 with ELIX UV/Progard 2) and dispensed from polyethylene water bottles.

Animals were fed a defined phytoestrogen-free diet (Product #D1010501, Research Diets, Inc. New Brunswick, NJ) either unsupplemented (control) or supplemented with BPA (2,2-bis(4- hydroxyphenyl)propane; CAS No. 80-05-7; Lot 11909; US EPA/NIEHS/NTP standard) or 17α- ethinyl estradiol (EE; 1,3,5(10)-estratrien-17α-ethinyl-3,17β-diol; CAS No. 57-63-6; Batch No. H923; Steraloids Inc., Newport, RI) that was homogenously incorporated into the diet by the manufacturer at desired final concentrations (BPA: 0.03 to 300 ppm which resulted in a dose range of 4 to 40,000 μg/kg/day; or EE: 0.0001 to 0.01 ppm, resulting in a dose range of 0.02 to 1.5 μg/kg/day ^22,24^. Food consumption was measured weekly and all mice were maintained on assigned dietary exposure from initiation of study, for the analyzed offspring (F1) this is at time of conception, until necropsy of offspring at ~12 weeks of age. Details of dietary consumption and resulting exposures to BPA and EE were reported previously ^22^.

Briefly, a total of 410 (205 each sex) Hsd:ICR mice (CD-1; Harlan Laboratories Indianapolis, IN) age six to seven weeks were mated (205 breeding pairs) following a 2 week acclimation period to assigned treatments. All treatment group and pairings assignments were done randomly upon arrival to the University of Cincinnati, Laboratory of Animal Science. After pairing, males were removed upon observation of a copulation plug or after two weeks without observed copulation. Females were then observed twice daily for any health-related outcomes, signs of pregnancy, and for parturition (designated postnatal day 0 or PND 0). Across all exposure groups there were no significant differences in length of gestation, litter size or sex ratio (Table 4). On PND1 the F1 offspring were assigned a coded study number and identified by a unique ear punch pattern. Offspring were separated from dams at PND 21, group housed 5 per cage by sex and maintained on dietary treatment until sacrifice. Litters were not culled and in cases of more than 5 males or 5 females per litter, same sex littermates were distributed evenly between two cages.

**Table 4.**
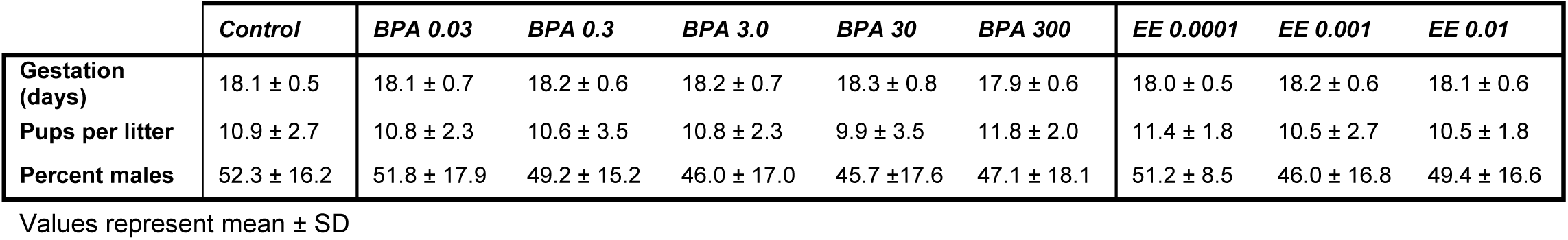
Litter Size and Sex Ratio

### Tissue preparation and histopathology

Tissue preparation and subsequent histopathologic analysis and scoring was done blinded to exposure group. Complete necropsy was performed for the F1 offspring at ~12 weeks of age (females PND 83 ± 6, range PND 74 - 97; males PND 82 ± 5, range PND 69 - 95). Females euthanized for necropsy were in the estrous stage of their cycle ^22^. Animals were euthanized by CO_2_ asphyxiation and perfused via cardiac puncture into the left ventricle with 10% neutral buffered formalin (Polysciences, Inc.; Warrington, PA), all tissues and organs were dissected and weighed. Tissues were then post-fixed with multiple changes of 10% neutral buffered formalin. The fixed tissues were blocked by sectioning with a sterile razor blade and washed several times in 70% ethanol prior to automated tissue processing and embedding in paraffin (Histocenter 3; Thermo-Shandon Kalamazoo, MI). Microtome sections were cut at 5 μm thickness from blocks at 4°C and placed on positively charged slides for standard hematoxylin and eosin (H&E) staining ^22^.

Microscopic examination of coded sections from the spleen of a single male and female from each litter was carried out separately by a board-certified pathologist (Diplomat of the American College of Veterinary Pathologists) following best-practices guidelines endorsed by the Society of Toxicologic Pathology and the European Society of Toxicologic Pathology for enhanced histopathology (EH) examination of the spleen from H&E stained slides ^32,34,57,58^. Briefly, the spleen was morphologically compartmentalized into white pulp and red pulp, and further compartmentalized into each of the microanatomic structures characteristic for the white pulp and red pulp compartment. Each compartment was assessed for changes in cellularity, resident and in-transit cell types, evidence of cell death (apoptosis or necrosis), accumulation of pigments (e.g. hemosiderin), and alterations in the extracellular matrix (i.e. fibrosis). Incidence and morphology for each microanatomical structure was scored using the recommended EH grading scheme of 0 = normal, 1 = minimal, 2 = mild, 3 = moderate, and 4 = marked ^32,36^.

### Statistical analysis

The statistical unit used was the litter for all analyses with males and females analyzed separately. All statistical analyses for differences in values compared to control were made independently for BPA and EE ^59^. Statistical analysis of data was performed using two-way or one-way analysis of variance (ANOVA), Dunnett’s multiple comparison post-hoc test, for ordinal histopathology data a rank order ANOVA Kruskal-Wallis H test with Dunn’s multiple comparisons tests, one-tailed χ^2^ test or a χ^2^ test for trend were used ^35,36^. GraphPad Prism v6 software (GraphPad Software, Inc., La Jolla, CA). A minimal level of statistical significance for differences in values among or between groups was considered p < .05.

## Acknowledgements

The authors are grateful for the outstanding histopathology analysis and guidance of Dr. Vinicius Carreira. This work was supported by the National Institutes of Health Grand Opportunity grant RC2ES018765 from the NIEHS that was awarded as part of the American Economic Recovery and Reinvestment Act, NIEHS grant R03ES023098, and by The NCSU Center for Health and Human Environment that is funded by NIEHS award P30ES025128.

## Author Contributions

R.G. performed the experiments, data collection, analysis, and contributed to writing of the manuscript. SB. conceived the experiments, performed data analysis and wrote the manuscript. Both authors have reviewed the manuscript.

